# PhenoMeNal: Processing and analysis of Metabolomics data in the Cloud

**DOI:** 10.1101/409151

**Authors:** Kristian Peters, James Bradbury, Sven Bergmann, Marco Capuccini, Marta Cascante, Pedro de Atauri, Timothy M D Ebbels, Carles Foguet, Robert Glen, Alejandra Gonzalez-Beltran, Ulrich Guenther, Evangelos Handakas, Thomas Hankemeier, Kenneth Haug, Stephanie Herman, Petr Holub, Massimiliano Izzo, Daniel Jacob, David Johnson, Fabien Jourdan, Namrata Kale, Ibrahim Karaman, Bita Khalili, Payam Emami Khonsari, Kim Kultima, Samuel Lampa, Anders Larsson, Christian Ludwig, Pablo Moreno, Steffen Neumann, Jon Ander Novella, Claire O’Donovan, Jake TM Pearce, Alina Peluso, Luca Pireddu, Marco Enrico Piras, Michelle AC Reed, Philippe Rocca-Serra, Pierrick Roger, Antonio Rosato, Rico Rueedi, Christoph Ruttkies, Noureddin Sadawi, Reza M Salek, Susanna-Assunta Sansone, Vitaly Selivanov, Ola Spjuth, Daniel Schober, Etienne A. Thévenot, Mattia Tomasoni, Merlijn van Rijswijk, Michael van Vliet, Mark R Viant, Ralf J. M. Weber, Gianluigi Zanetti, Christoph Steinbeck

**Affiliations:** Leibniz Institute of Plant Biochemistry, Stress and Developmental Biology, Weinberg 3, 06120 Halle (Saale), Germany; School of Biosciences, University of Birmingham, Edgbaston, Birmingham, B15 2TT, United Kingdom; Department of Computational Biology, University of Lausanne, Lausanne, Switzerland; Swiss Institute of Bioinformatics, Lausanne, Switzerland; Division of Scientific Computing, Department of Information Technology, Uppsala University, Sweden; Department of Pharmaceutical Biosciences, Uppsala University, Box 591, 751 24 Uppsala, Sweden; Department of Biochemistry and Molecular Biomedicine, Universitat de Barcelona; Centro de Investigación Biomédica en Red de Enfermedades Hepáticas y Digestivas; Department of Biochemistry and Molecular Biomedicine, Universitat de Barcelona; Centro de Investigación Biomédica en Red de Enfermedades Hepáticas y Digestivas (CIBEREHD), Instituto de Salud Carlos III (ISCIII), Spain; Department of Surgery & Cancer, Imperial College London, South Kensington, London, SW7 2AZ, United Kingdom; Centre for Molecular Informatics, Department of Chemistry, University of Cambridge, Lensfield Road, Cambridge, CB21EW, United Kingdom; Oxford e-Research Centre, Department of Engineering Science, University of Oxford, 7 Keble Road, OX1 3QG, Oxford, UK.; Netherlands Metabolomics Center, Leiden, 2333 CC, Netherlands; Division of Systems Biomedicine and Pharmacology, Leiden Academic Centre for Drug; European Molecular Biology Laboratory, European Bioinformatics Institute (EMBL-EBI), Wellcome Genome Campus, Hinxton, Cambridge CB10 1SD, UK; Department of Medical Sciences, Clinical Chemistry, Uppsala University, 751 85 Uppsala, Sweden; INRA, University of Bordeaux, Plateforme Métabolome Bordeaux-MetaboHUB, 33140 Villenave d’Ornon, France; INRA - French National Institute for Agricultural Research, UMR1331, Toxalim, Research Centre in Food Toxicology, Toulouse, France; Department of Epidemiology and Biostatistics, School of Public Health, Imperial College London, St. Mary’s Campus, Norfolk Place, W2 1PG, London, United Kingdom; National Bioinformatics Infrastructure Sweden, Uppsala University, Uppsala, Sweden Department of Pharmaceutical Biosciences, Uppsala University, Uppsala, Sweden; German Centre for Integrative Biodiversity Research (iDiv) Halle-Jena-Leipzig, Deutscher Platz 5e, 04103 Leipzig, Germany; Distributed Computing Group, CRS4, Pula, Italy; College of Medical and Dental Sciences, University of Birmingham, Edgbaston, Birmingham, B15 2TT, United Kingdom; CEA, LIST, Laboratory for Data Analysis and Systems’ Intelligence, MetaboHUB, Gif-Sur-Yvette F-91191, France; Magnetic Resonance Center (CERM) and Department of Chemistry, University of Florence and CIRMMP, 50019 Sesto Fiorentino, Florence, Italy; European Bioinformatics Institute (EMBL-EBI), European Molecular Biology Laboratory, Wellcome Genome Campus, Hinxton, Cambridge CB10 1SD, U.K; ELIXIR-NL, Dutch Techcentre for Life Sciences, Utrecht, 3503 RM, Netherlands; Netherlands Metabolomics Center, Leiden, 2333 CC, The Netherlands; Division of Systems Biomedicine and Pharmacology, Leiden Academic Centre for Drug Research (LACDR), Leiden University, Leiden, 2333 CC, The Netherlands; BBMRI-ERIC, Graz, Austria; Cheminformatics and Computational Metabolomics, Institute for Analytical Chemistry, Lessingstr. 8, 07743 Jena, Germany

**Keywords:** Metabolomics, Data Analysis, e-infrastructures, NMR, Mass Spectrometry, Computational Workflows, Galaxy, Cloud Computing, Standardization, Statistics

## Abstract

**Background:** Metabolomics is the comprehensive study of a multitude of small molecules to gain insight into an organism’s metabolism. The research field is dynamic and expanding with applications across biomedical, biotechnological and many other applied biological domains. Its computationally-intensive nature has driven requirements for open data formats, data repositories and data analysis tools. However, the rapid progress has resulted in a mosaic of independent – and sometimes incompatible – analysis methods that are difficult to connect into a useful and complete data analysis solution.

**Findings:** The PhenoMeNal (Phenome and Metabolome aNalysis) e-infrastructure provides a complete, workflow-oriented, interoperable metabolomics data analysis solution for a modern infrastructure-as-a-service (IaaS) cloud platform. PhenoMeNal seamlessly integrates a wide array of existing open source tools which are tested and packaged as Docker containers through the project’s continuous integration process and deployed based on a kubernetes orchestration framework. It also provides a number of standardized, automated and published analysis workflows in the user interfaces Galaxy, Jupyter, Luigi and Pachyderm.

**Conclusions:** PhenoMeNal constitutes a keystone solution in cloud infrastructures available for metabolomics. It provides scientists with a ready-to-use, workflow-driven, reproducible and shareable data analysis platform harmonizing the software installation and configuration through user-friendly web interfaces. The deployed cloud environments can be dynamically scaled to enable large-scale analyses which are interfaced through standard data formats, versioned, and have been tested for reproducibility and interoperability. The flexible implementation of PhenoMeNal allows easy adaptation of the infrastructure to other application areas and ‘omics research domains.

## Findings

## Background

The field of metabolomics has seen remarkable progress over the last decade and has enabled fascinating discoveries in many different research areas. Metabolomics is the study of small molecules in organisms which can reveal detailed insights into metabolic biochemistry, e.g. changes in concentrations of specific molecules, metabolic fluxes between cells or compartments, identification of molecules that are involved in the pathogenesis of a disease, the study of the biochemical phenotype of animals, plants and even soil microorganisms [1–3]

The principal metabolomics technologies of mass spectrometry (MS) and nuclear magnetic resonance spectroscopy (NMR) typically generate large data sets that require computationally intensive analyses [4]. For example, biomedical investigations can involve large cohorts with many thousands of metabolite profiles and can produce terabytes of data [5]. With such large data sets, processing becomes impracticable and unmanageable on commodity hardware. Cloud computing can offer a solution by enabling the outsourcing of calculations from local workstations to scalable cloud data centers, with the possibility to allocate thousands of CPU cores simultaneously. Furthermore, cloud computing allows for resources to be instantiated on-demand (CPUs, RAM, network, storage) and access to computational tools in the form of microservices that can dynamically grow or shrink.

MS and NMR data processing usually involves selection of parameters (which are often specific to the analytical instrumentation), algorithmic peak detection, peak alignment and grouping, annotation of putative compounds and extensive statistical analyses [6,7]. Many open source tools have been developed that address these different steps in data processing and analysis. These tools, however, usually come with their own software dependencies, resource requirements and scripting languages. As a consequence, configuring and running them is often complicated, especially for researchers who are untrained in computer science [4]. Furthermore, many tools require users to input parameters that can significantly affect results and performance, and reporting of these parameters is not always clear [8].

In the last five years, a number of infrastructures and integration efforts were initiated, including metabolomics data repositories with a global scope [9,10], platforms for reproducible workflow analysis [11,12], as well as initiatives to integrate and coordinate data standards [13]. Simultaneously, multiple networks of service centers such as the international Phenome Centers and MetaboHub (http://phenomenetwork.org, http://www.metabohub.fr/metabohub.html) have formed with the goal to facilitate the acquisition, processing and analysis of metabolomics data [9,14,15] at ever increasing scales.

Here we present PhenoMeNal, a robust and performant data analysis e-infrastructure that provides a large suite of standardized and interoperable metabolomics data processing tools as a complete data analysis solution. The PhenoMeNal e-infrastructure can be easily deployed onto public and private cloud environments, enabling scalable and cost-effective high-performance metabolomics data analysis in a way that hides the technical complexity from the user. PhenoMeNal facilitates reproducible analyses through automated, sharable and citable workflows.

## Overview

The features of the PhenoMeNal e-infrastructure are encapsulated as a Cloud Research Environment (CRE). The PhenoMeNal CRE can be instantiated on major commercial public cloud providers, including Amazon Web Services (AWS) and Google Cloud Platform (GCE), as well as OpenStack-based private clouds. Technical complexity is hidden from the users, simplifying setting up the cloud infrastructure for administrators. From a web-based portal, users can deploy the CRE, which includes several web services and software tools. Data can be processed directly in the e-infrastructure without the need to install additional software. Scientific workflows can be executed via user friendly web-based platforms such as Galaxy, as well as programmatic interfaces and notebooks. Each service has been supplied with a rich source of documentation and training material to assist researchers.

**Fig. 1:**
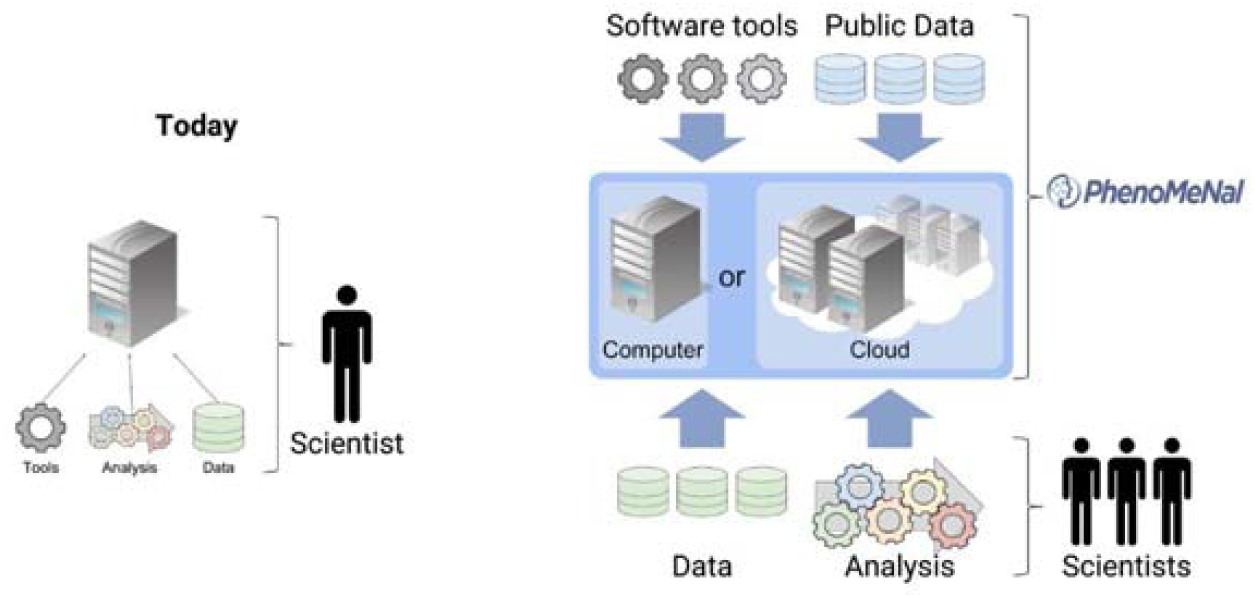
Conceptual design of the PhenoMeNal cloud e-infrastructure (right hand side) compared to traditional approaches in biomedicine (left side).

### The PhenoMeNal Portal

The PhenoMeNal portal (https://portal.phenomenal-h2020.eu/) allows users to deploy, manage and delete PhenoMeNal CREs simply through a web interface. Deployments to the three major commercial cloud platforms (AWS and GCP) as well as OpenStack, an open source cloud platform, can be made using an easy to follow wizard (Fig. 2). OpenStack deployments can be deployed behind clinical firewalls, which is especially pertinent when dealing with sensitive (i.e. patient) data.

**Fig. 2:**
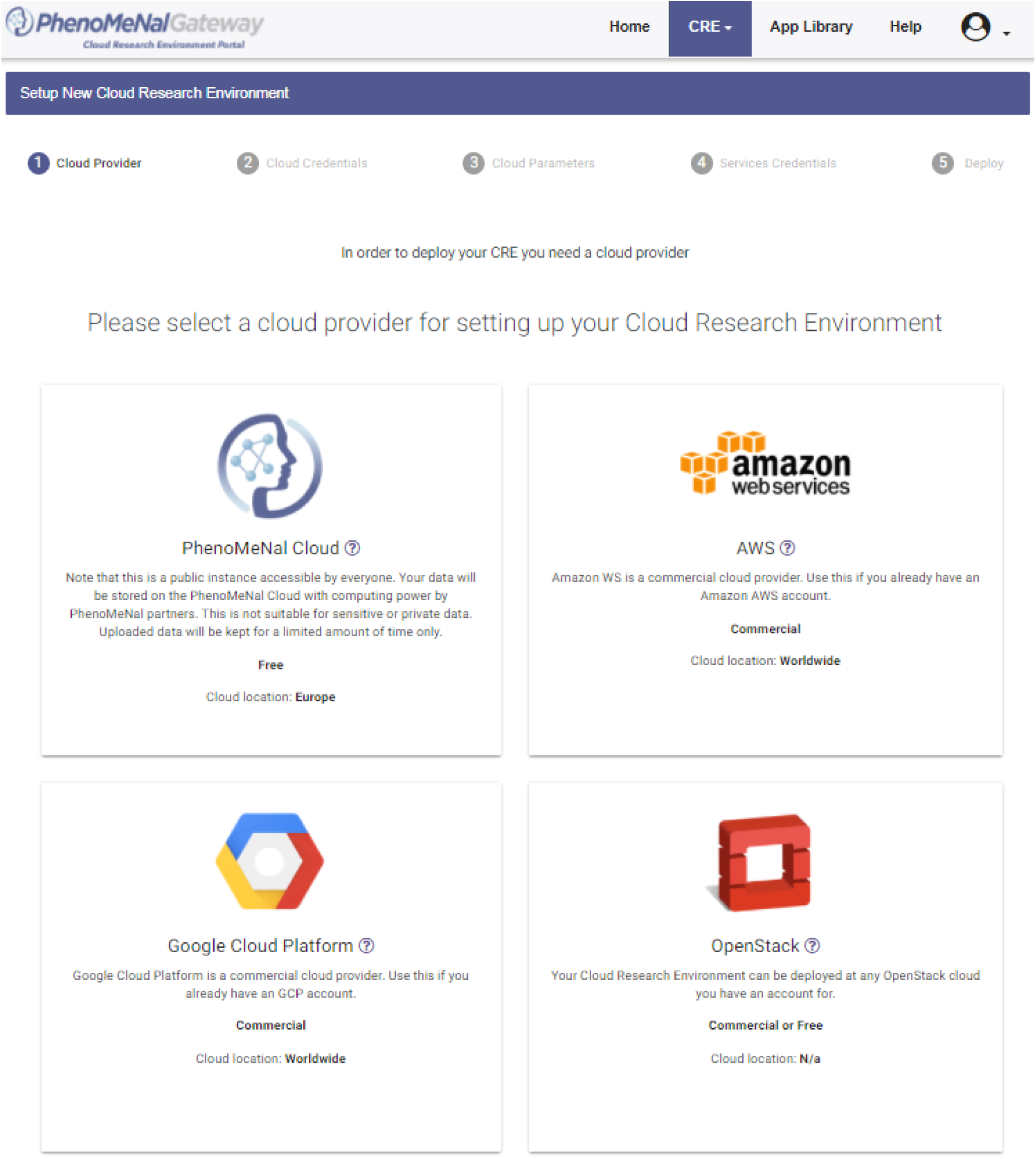
Screenshot of selecting a cloud provider during setup of the Cloud Research Environment (CRE) using the dedicated PhenoMeNal portal.

The PhenoMeNal public instance allows users to test-run a CRE without the need to deploy on a cloud platform. It can be deployed and accessed through the portal (Fig. 2). Once credentials for users have been generated, analyses can be run through a Galaxy instance containing the tools and workflows present in any deployed CRE. The portal also includes user and developer documentation, workflow tutorials and links to training videos.

### Scientific workflows

A scientific workflow is a set of computational steps that are carried out to process and analyze data [16]. Usually, a workflow is comprised of several linked software tools that are each executed during a particular step of the workflow. In order to manage and automate scientific workflows, PhenoMeNal uses the well-established dedicated workflow management system Galaxy, which presents the user with an easy to use graphical user interface as well as providing a programmatic interface [17,18]. Galaxy facilitates collaborative exchange, reproducibility and traceability of data analysis by enabling users to share entire workflows and analysis histories [19]. In addition to Galaxy, programmatic executable notebooks (Jupyter) and the programmatic interfaces Luigi and Pachyderm are also supported [20].

In order to cover typical use cases in metabolomics and to illustrate the usage and applicability of given analytical pipelines and software tools, five representative scientific workflows are available in the PhenoMeNal CRE Galaxy environment (Table 2, Fig. 6), each having different computational demands and purposes. More than 250 individual modules have been integrated in Galaxy (see section scientific workflows in Methods).

**Table 2:** List of workflows which are representative for their respective metabolomics domains (Identification in NMR, Fluxomics, Annotation and identification in MS and eco-metabolomics).

| Workflow name | Description | References |
| --- | --- | --- |
| 1D NMR | Processes 1D NMR experiments from raw data to a data matrix required for visualisation and statistical analysis, building on nmrML and NMRProcFlow. The automatic workflow is based on the MTBLS1 data set, describing urinary changes in type 2 diabetes in humans. | [30,70,71] |
| Fluxomics | Quantifies steady state fluxes following <sup>13</sup>C Metabolic Flux Analysis. The workflow was first based on the analysis of the MTBLS412 data set with <sup>13</sup>C tracer data of human umbilical vein endothelial cells (HUVEC) under hypoxia. | [72,73] |
| LC-MS/MS | Processes, quantifies and annotates/identifies features in mass spectra using MetFrag - a tool which annotates molecules from compound databases of tandem mass spectrometry (MS/MS) spectra. The workflow is based on MTBLS558. | [55,69,74] |
| Univariate and Multivariate Statistics | Applies univariate and multivariate statistical analysis, and illustrates how data sets may be explored enabling the identification of variables of interest and the construction of predictive models. The workflow is based on MTBLS404. | [11,38] |
| Eco-Metabolomics | Implementation of a resource demanding metabolomics use case in ecology, used in large field experiments to describe interactions between different species of organisms in remarkable detail. The workflow is based on MTBLS520. | [44] |
| ISA-Create-Validate-Upload | A workflow to create ISA compliant metadata files based on study design information, augmented with semantic markup as source, implementing UK Phenome center naming conventions. Following validation, the workflow also allows visualization of overall study design and deposition to EMBL-EBI |  |

**Fig. 6:**
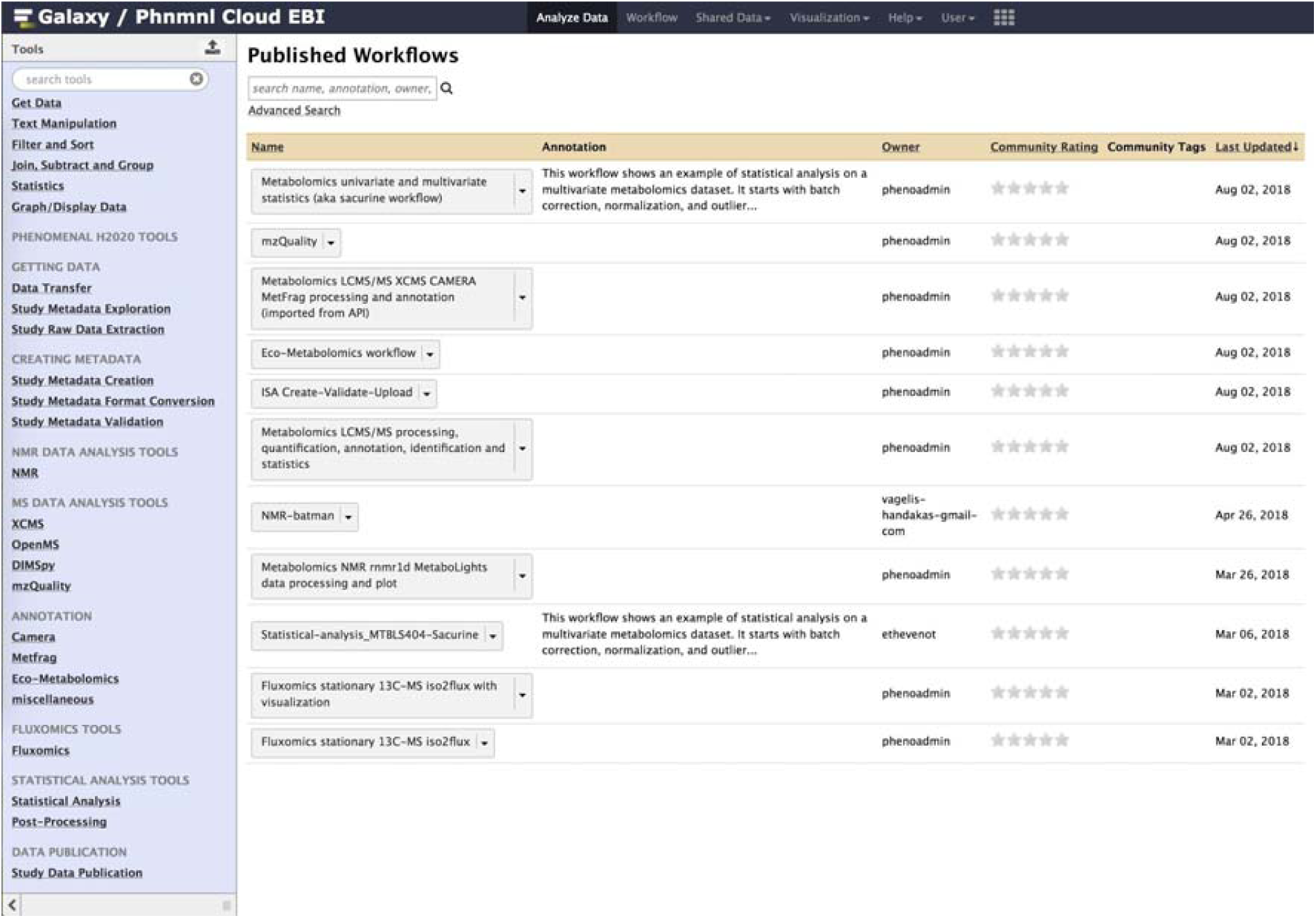
Screenshot of the available workflows in Galaxy.

### Software tools

The Portal App Library (https://portal.phenomenal-h2020.eu/app-library) shows all the software tools packaged in PhenoMeNal that are available through the CRE deployment (Fig. 3). The range of software tools available cover several metabolomics domains, making PhenoMeNal relevant for use in a wide range of data analysis scenarios. The domains covered include clinical metabolomics, plant metabolomics, fluxomics and eco-metabolomics. Data from both targeted and untargeted analysis can be analyzed for metabolite profiling and fingerprinting approaches [1,2]. NMR and MS (LC/MS, GC/MS, DIMS) data can be processed.

**Fig. 3:**
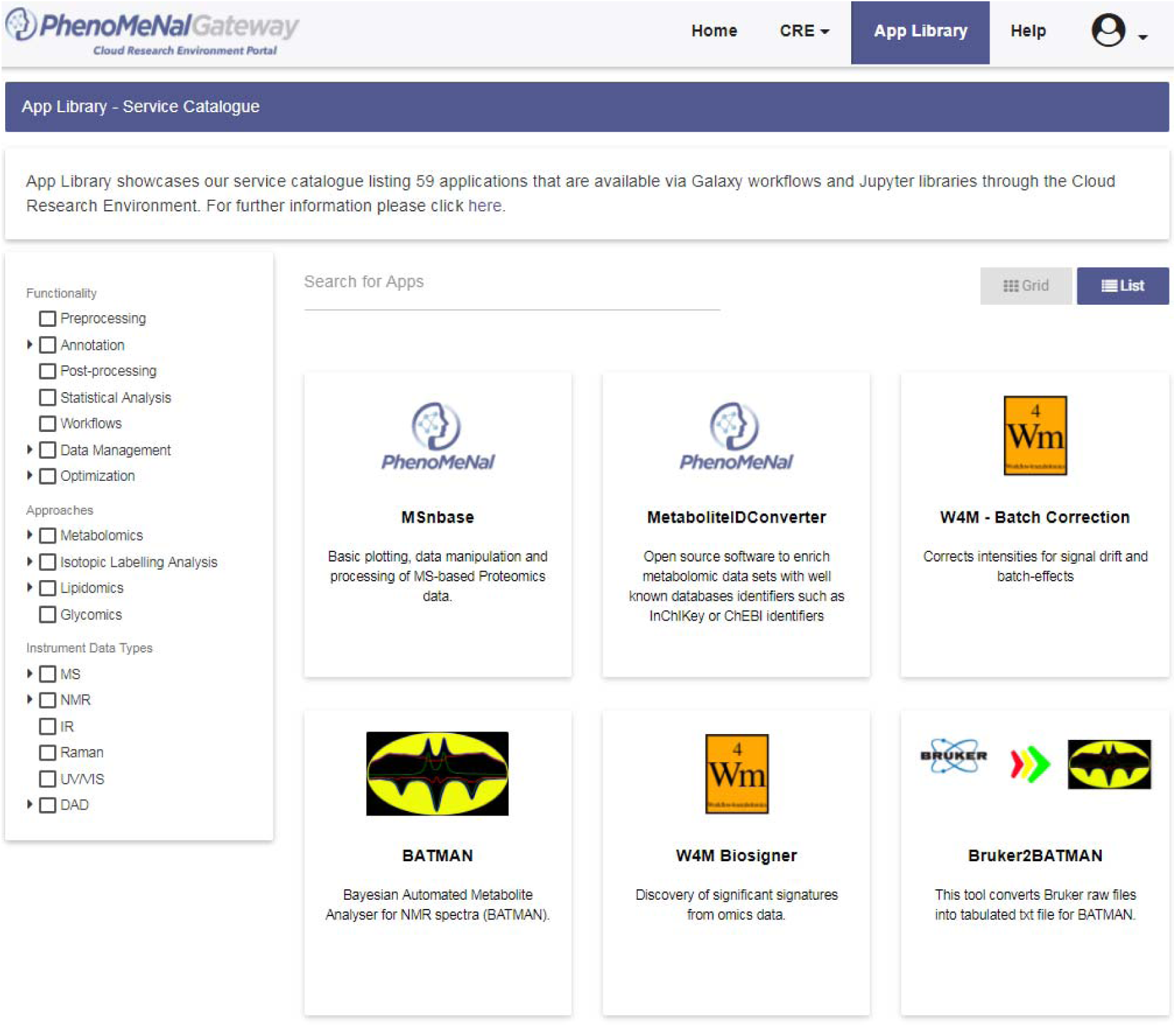
Screenshot of the PhenoMeNal Portal App library.

PhenoMeNal also provides tools for data management (e.g. via the ISA format and API), metabolite feature detection (e.g. XCMS, CAMERA, nmrProcFlow), metabolite identification (MetFrag, BATMAN, MetaboMatching) and (bio)statistics (e.g. univariate, multivariate and power analyses) (Table 1). Tools can be filtered for functionality, approaches and instrument (data) types to readily find the most appropriate software tools (Fig. 3). Some tools that implement specific functionality (e.g. Rnmr1D which performs baseline correction of NMR spectra as part of nmrProcFlow) are available through dedicated Galaxy modules or through software containers (Table 1).

**Table 1:**
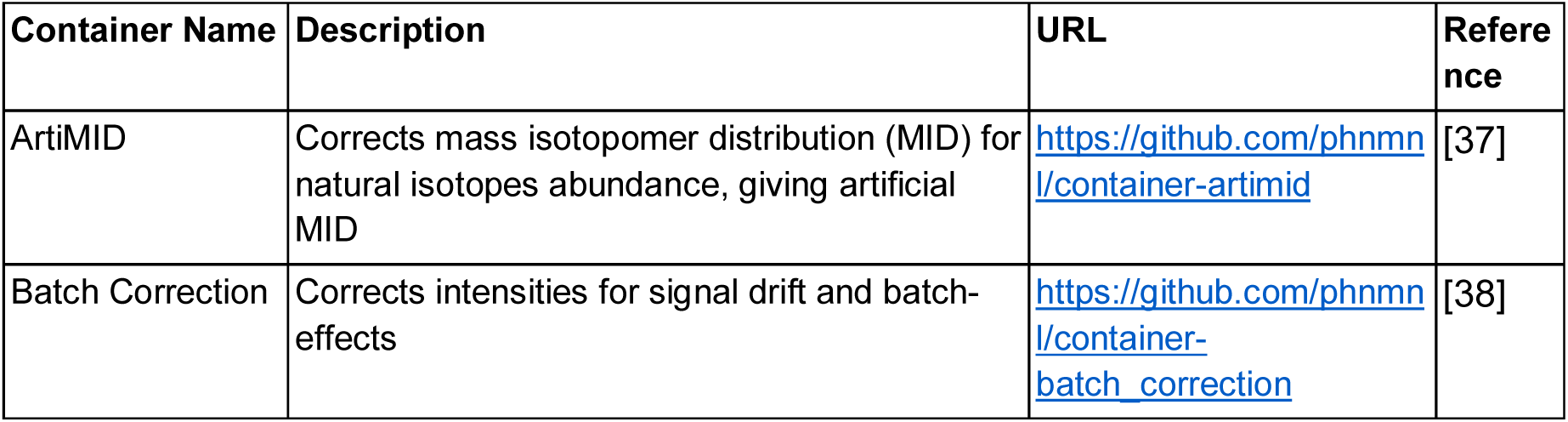
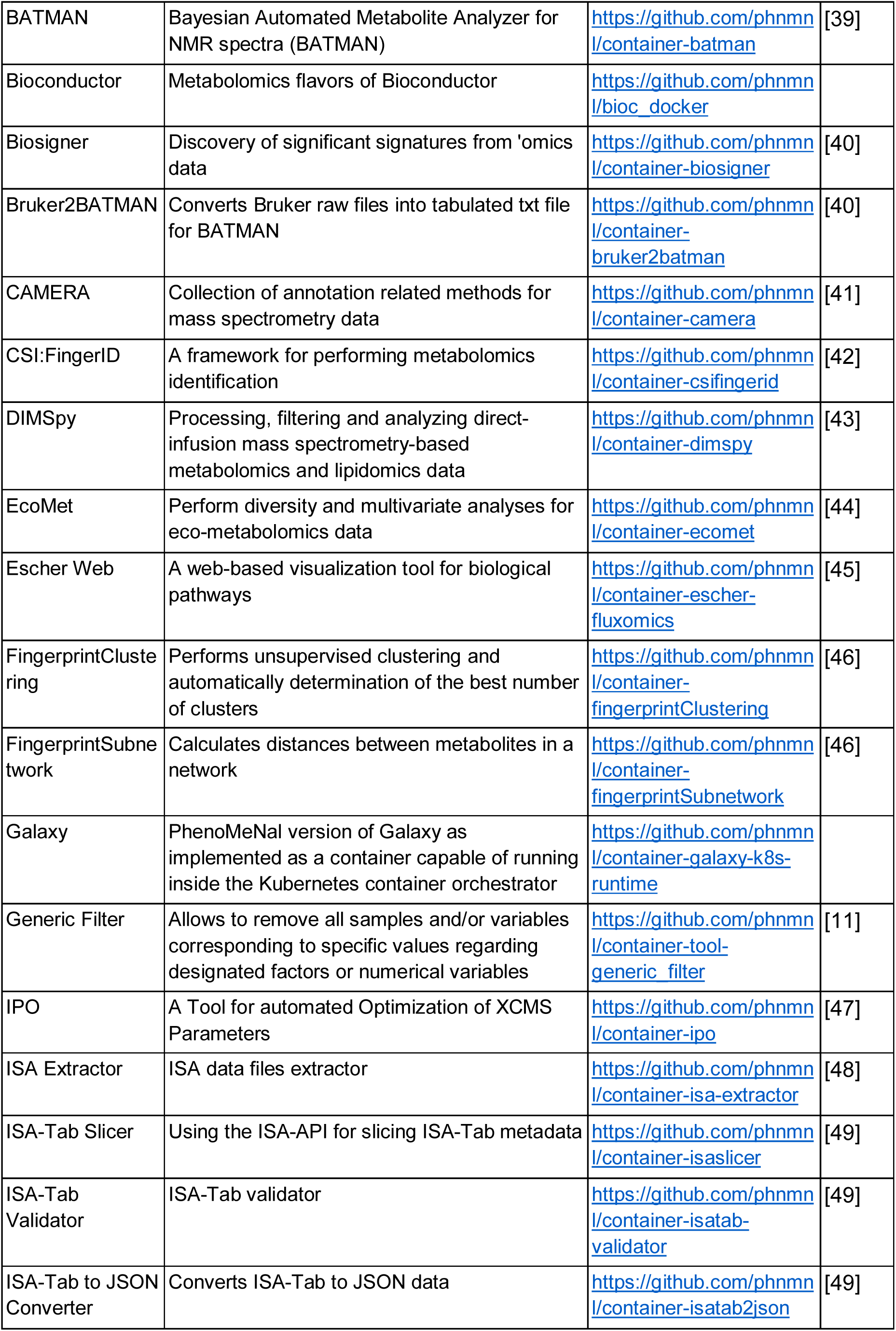
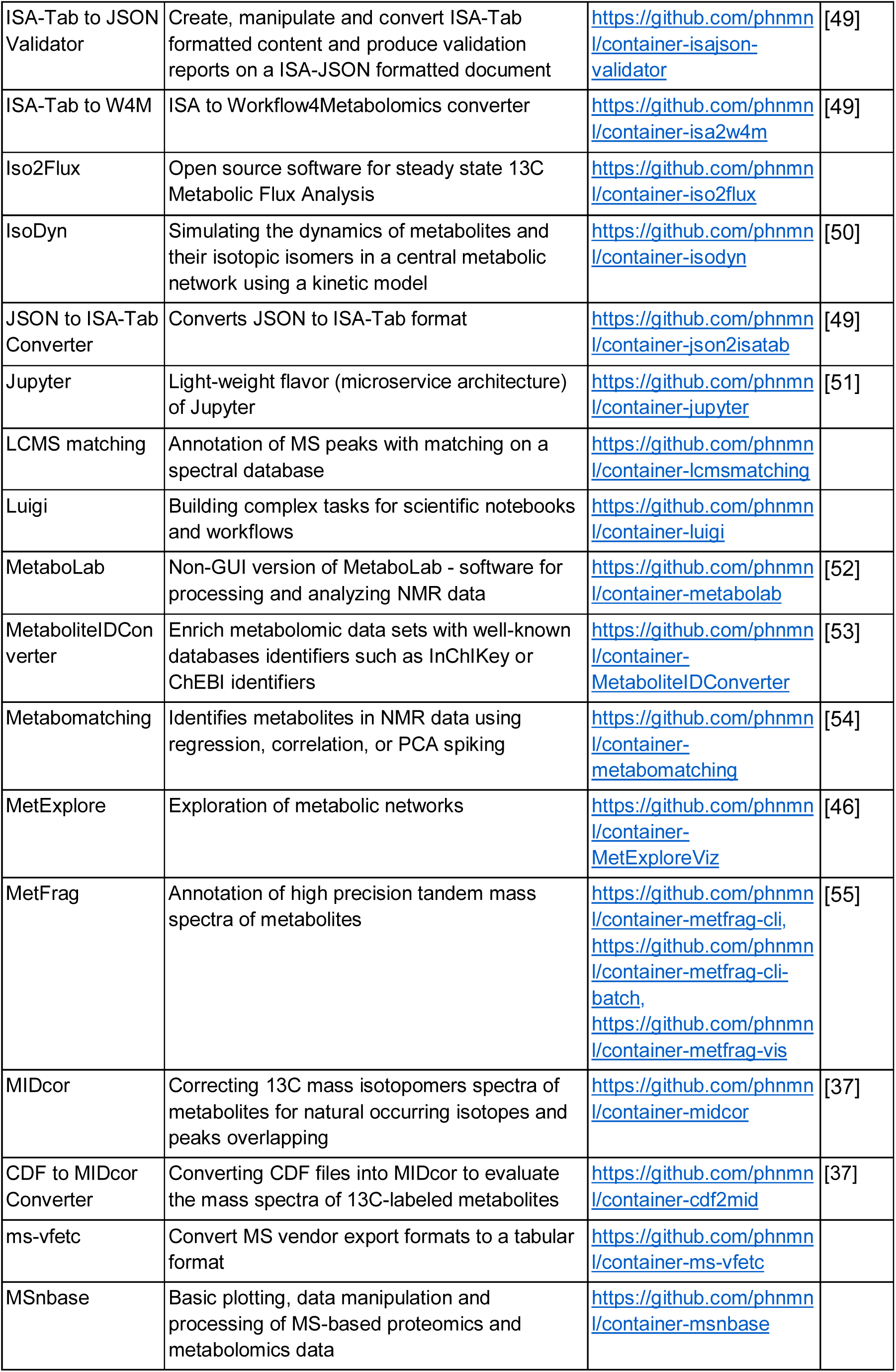
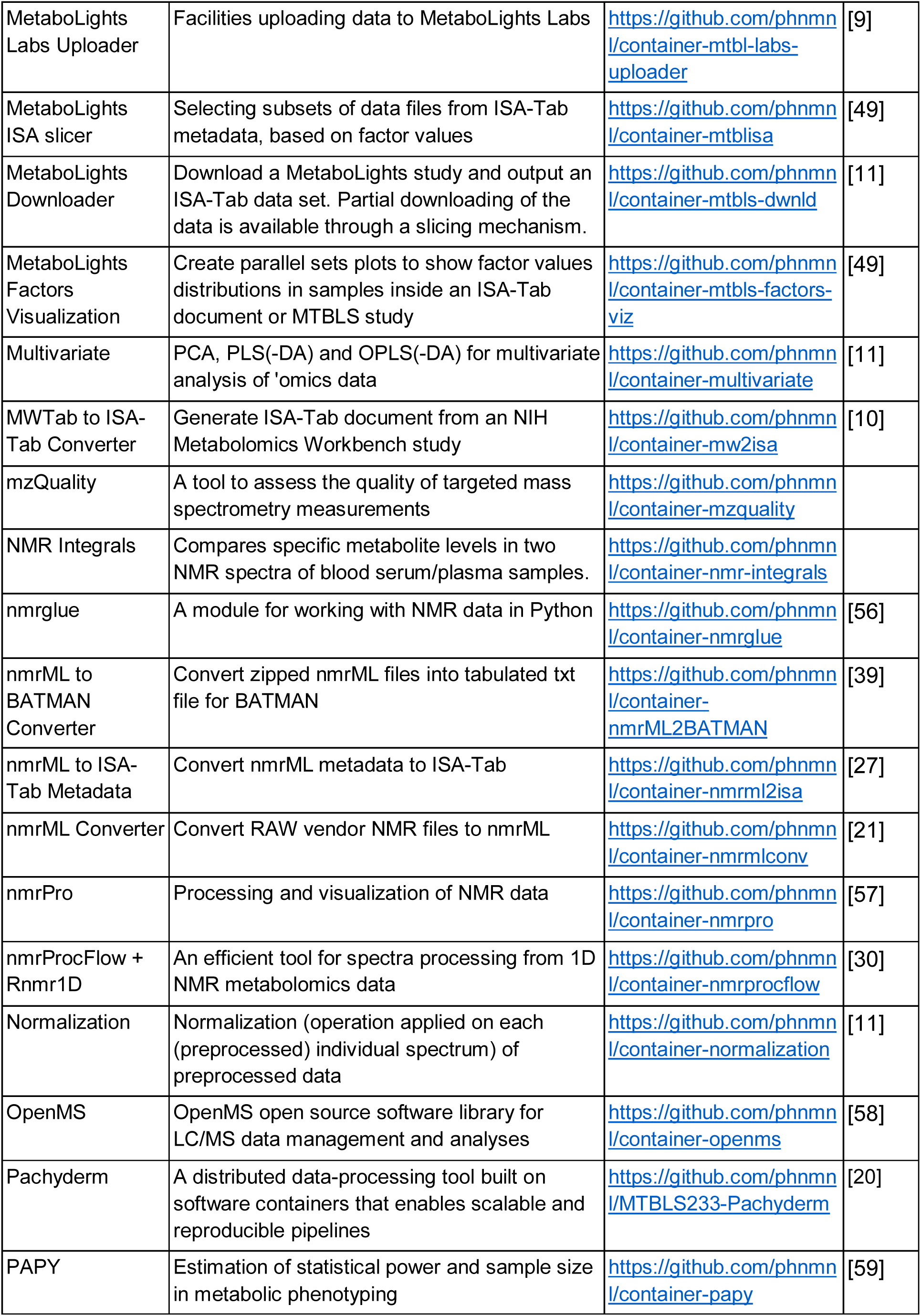
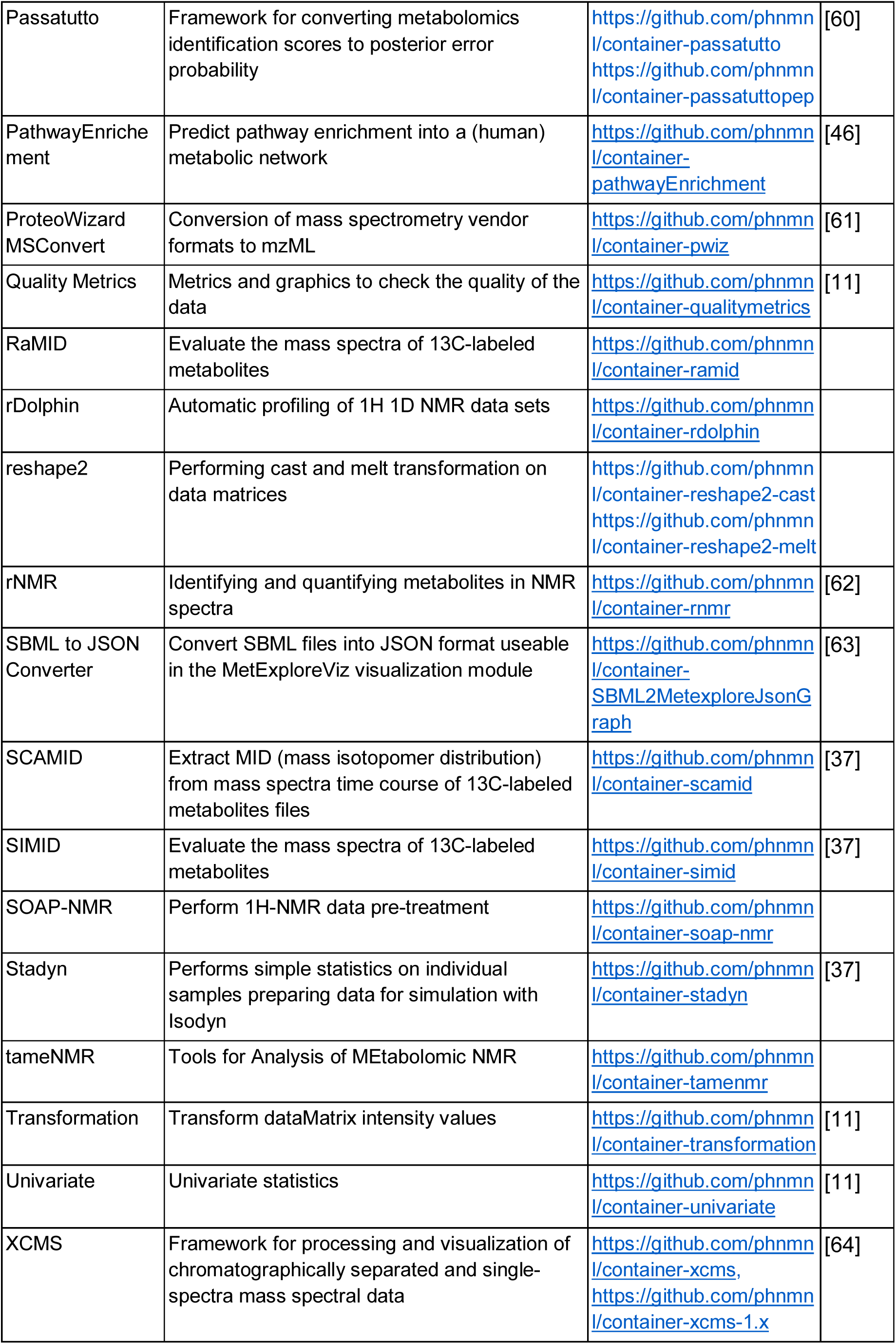
List of external software tools that were incorporated into PhenoMeNal.

### Study Design

PhenoMeNal was designed to use standardized protocols, software tools and comply with state-of-the-art dedicated specifications and data formats across the entire project. Development was geared towards implementation of open standards for tracking provenance of both data and metadata generated by clinical phenotyping projects. In PhenoMeNal, the ISA model and specifications were implemented using the ISA format to generate, annotate, validate and deposit experimental metadata information of data sets and studies to public repositories such as MetaboLights [21,22]. ISA based metadata tracking is used for the different analysis pipelines which are specific to the distinct metabolomics domains. PhenoMeNal reached native support for the ISA format by developing a dedicated Galaxy composite data type. Such component affords direct recognition of the ISA format by the Galaxy environment, thus ensuring seamless integration with downstream workflow component.

### Data deposition

PhenoMeNal encourages the metabolomics data repository MetaboLights as a primary source of data deposition [23]. Private and public data sets are supported, as well as download and upload to MetaboLights. If the storage in a data repository such as MetaboLights is not possible, data can be stored locally or in the cloud e-infrastructure. Access to the data is strictly controlled and secured. To support data deposition, ISA based Galaxy modules are available allowing to publish and disseminate scientific results in standard compliant ways (see also Table 2).

### Reproducibility

One of the challenges of cloud computing is that analyses need to be run continuously and successfully in different environments [24]. Specifically, it has to be ensured that, given the same input, workflows and tools produce identical results regardless of the underlying environment [4,24]. When these requirements are fulfilled, end users can be confident that their data will be analyzed correctly. PhenoMeNal has implemented three major testing strategies to ensure technical reproducibility using a continuous integration framework [25]. Tests were implemented for the infrastructure components, individual software containers and for data involved in computational workflows.

### Sustainability

PhenoMeNal is part of a number of initiatives (BioMedBridges, COSMOS and ELIXIR) to foster the role of metabolomics and to harmonize experimental data and metadata usage [13,26]. Collaborations were established with EGI and Indigo Datacloud, to ensure that PhenoMeNal uses technologies that are well-supported and assure their widespread usage, continuity and further development. For example, the development of KubeNow and contributions to the Galaxy and Workflow4Metabolomics community are essential for PhenoMeNal. Core development will continue on GitHub and is fostered by collaborations with tool developers.

Dependencies on specific technologies and frameworks were avoided by focusing on open standards such as ISA-Tab / ISA-JSON, mzML and nmrML and widely accepted software [27]. By being able to deploy PhenoMeNal on multiple types of cloud environments, lock-in to specific computing resource providers are avoided. PhenoMeNal implemented continuous integration and delivery, validated by extensive testing and with clear maintenance responsibilities.

### Privacy and security

With human or animal material the collection, storage and analysis of metabolomics data introduce a number of constraints due to Ethical, Legal and Social Implications (ELSI) [28]. In particular, data initially derived from clinical studies may be identifiable and will require consent for use, usually for a defined objective such as diagnosis, or be related to a particular disease study. PhenoMeNal has support for fully anonymized and pseudonymized data (where identifiers are removed, but data can be identified through a mapping such as a hash or code) [29]. In general, PhenoMeNal implements and is fully compliant to the ELSI guidelines [28]. PhenoMeNal has been designed to accommodate cases where sensitive data needs to be processed according to the General Data Protection Regulation (GDPR).

With sensitive or private data, users can always deploy PhenoMeNal on local resources only, thus avoiding the export of data altogether. Access to the e-infrastructure is strictly controlled through authorization. Users must register in order to use the individual parts of the e-infrastructure. Transport and network communications are secured and encrypted. PhenoMeNal is part of the NeIC-Tryggve2 (https://neic.no/tryggve2/) project to further steer the development of secure data storage of biomedical research data.

### Documentation and Training materials

Extensive user documentation and tutorials are provided via the PhenoMeNal Wiki page (https://github.com/phnmnl/phenomenal-h2020/wiki). The Wiki includes detailed developer resources including information about the PhenoMeNal release schedule, guidelines for tool, workflow and portal developers, continuous integration and testing. Further documentation detailing, creating and managing PhenoMeNal CREs, tutorials for the Galaxy modules and pre-configured workflows, as well as Galaxy tours that provide step by step guidance for inexperienced users are also provided.

### Community engagement

The PhenoMeNal project is open source, and is hosted on GitHub (https://github.com/phnmnl/). Developers can contribute tools to PhenoMeNal and are encouraged to do so. To add a tool to PhenoMeNal, it must be containerized using Docker, and then integrated into the build process. Detailed documentation is available in the project’s Wiki for developers who wish to add their tools to the PhenoMeNal CRE.

Collaborations with other projects have been actively encouraged during the development of PhenoMeNal, including Workflow4Metabolomics [11] and the developers of both nmrML and nmrProcFlow [30]. These collaborations are essential to foster greater standardization within PhenoMeNal and to increase compatibility with other metabolomics data processing infrastructures.

## Availability

Information on how to access PhenoMeNal can be found at https://phenomenal-h2020.eu. The GitHub repository https://github.com/phnmnl/ hosts the source code of all development projects. The project container-galaxy-k8s-runtime contains all of the developments regarding Galaxy. The Wiki containing documentation is also hosted on GitHub https://github.com/phnmnl/phenomenal-h2020/wiki. The PhenoMeNal Portal can be reached at https://portal.phenomenal-h2020.eu. The public instance of Galaxy is accessible at https://public.phenomenal-h2020.eu. Source code and documentation are available under the terms of the Apache 2.0 license. Integrated open source projects are available under the respective licensing terms.

## Conclusions

PhenoMeNal has succeeded in increasing the robustness and coverage of representative metabolomics data sets and workflows. Our efforts were also guided by feedback from real life test scenarios collected at workshops with users from the clinical domain. The e-infrastructure covers a wide range of analysis pipelines including data generation and download, data pre- and post-processing, (bio)statistics and result deposition in appropriate data repositories. PhenoMeNal has fostered the visibility of new metabolomics tools and has enabled the development of more sophisticated data analysis workflows. A large effort has been made to introduce lower level changes to cloud e-infrastructures (e.g. the cloud deployment software KubeNow) to meet the demands of the biomedical domain. Furthermore, Galaxy has been enriched with metabolomics data standards, in particular the ISA format for study metadata and mzML and nmrML for acquired data files, as well as support for Kubernetes.

PhenoMeNal constitutes a keystone solution in cloud platforms available for metabolomics data analysis. The platform was designed to deliver optimal performance and functionality for typical use cases in the metabolomics domain. While the needs of clinicians and researchers in the biomedical and biochemical domains have been targeted, PhenoMeNal is not limited to a specific domain as the cloud infrastructure, tools and workflows can be adapted to other use cases as demonstrated with the inclusion of the eco-metabolomics workflow. The technological advancements can be reused in other scientific cloud environments and could be integrated with solutions from other ‘omics domains in the future.

## Methods

### Cloud e-infrastructure

The PhenoMeNal CRE is designed as a microservice architecture, with services being implemented as Virtual Machine Images (VMIs) and software containers. Containers are used to provision microservices for metabolomics data analysis tools and also long-running services such as workflow management systems. A container orchestrator runs containers on top of the scalable infrastructure. The orchestrator takes a group of machines that act as a distributed cluster and receives requests for tools as well as services executions. PhenoMeNal implements various layers to provision a container orchestrator on top of either bare metal hardware or Infrastructure as a Service (IaaS) given by a cloud provider [31] (Fig. 4).

**Fig. 4:**
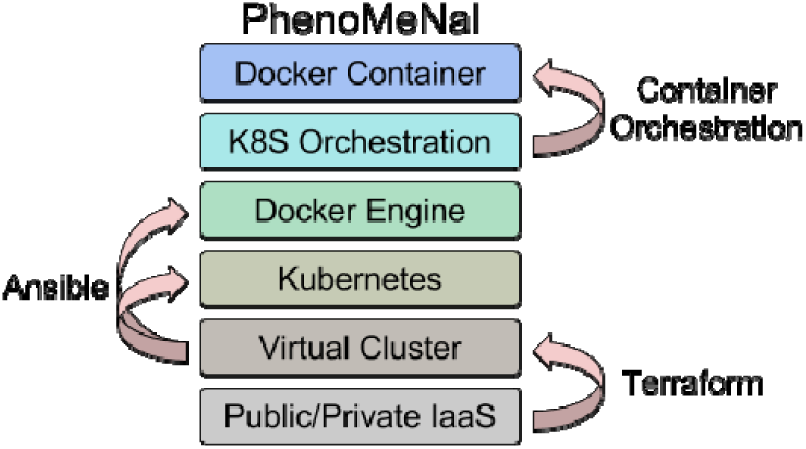
PhenoMeNal implements various layers to provision containers on top of the e-infrastructure.

Starting from the IaaS, the first layer is a cluster of Virtual Machines (VMs) which are started and initialized with a defined operating system (from a base image). This is called infrastructure provisioning, and in PhenoMeNal VMs are executed through the Terraform framework [32]. Terraform deploys VM setups to a number of public and private cloud providers including OpenStack, GCE and AWS. The resulting VMs run with a clean install of an operating system including the relevant networking features. Google Kubernetes is used to run software on top of the provisioned VMs [20]]. The Ansible framework is used for the software provisioning layer which performs the deployment of the container daemon and the container orchestrator [33]. Docker is used as the orchestrator daemon for the containers [34].

PhenoMeNal provides IaaS for three different cloud environments:

1. “local cloud”: local workstations or clusters where data are not allowed to leave the facility.
2. “public cloud”: the flexible use of commercial cloud providers such as GCP and AWS.
3. “shared cloud”: using OpenStack - a free and open-source software platform for cloud computing, ideal for custom environments and research networks.

The cloud infrastructure of PhenoMeNal is based upon containers that are deployed in a Kubernetes environment. Deployment is managed by KubeNow, which is developed by the PhenoMeNal team in order to simplify managing the deployment, including storage, network and other required services [20,35]. Orchestration is handled by using Helm charts. The storage subsystem is based on the cloud storage file system GlusterFS. Security is guaranteed via HTTPS encryption (SSL certificates issued by Cloudflare). This elastic implementation allows PhenoMeNal to be instantiated on any Kubernetes-based cloud environment [16]. We use a standardized REST API to operate and communicate between the different interfaces [36].

### Software tools

The PhenoMeNal portal has an Application Library that allows users to deploy tools as microservices into the cloud infrastructure (Table 1, Fig. 5). The portal is packaged into frontend and backend engines on top of Kubernetes. Orchestration is managed by Helm charts.

**Fig. 5:**
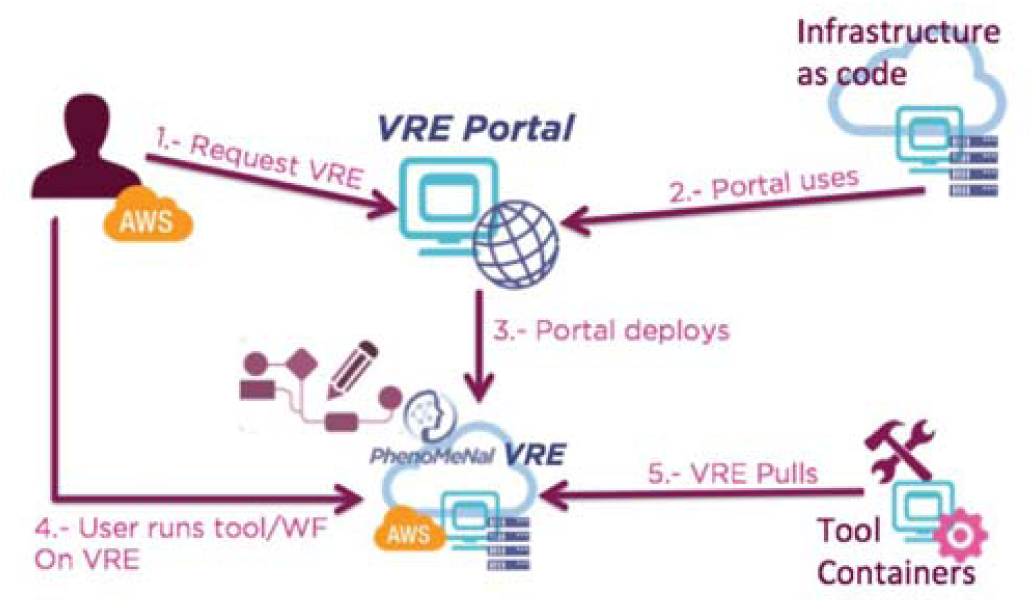
User interaction with the central PhenoMeNal components. Starting from the CRE Portal users can deploy infrastructure components and services choosing from different cloud providers. When the deployment has been made, services can be used privately.

Most software tools in PhenoMeNal are compiled from source code and use a variety of programming languages. Linux versions of software tools and user interfaces such as Galaxy are supported in dedicated encapsulated Docker containers which are implemented as minimum-sized microservices. PhenoMeNal currently hosts 100 such projects in its GitHub repository (https://github.com/phnmnl/?q=container). Projects are indicated by the trailing ‘container-’ name and include a ruleset to build and run the containerized tools, as well as data sets for testing and other necessary files.

### Scientific workflows

The Galaxy workflow management system has continued to develop for nearly a decade and is widely regarded as one of the most popular scientific workflow platforms [17,65]. It provides a user-friendly web-based graphical user interface to make it easy for the end-user to configure and run individual modules and entire workflows without programming experience. Galaxy wraps command-line tools and scripts into modules that are launched via the web interface. In PhenoMeNal, we encapsulated each Galaxy module into a Docker container that can be flexibly launched in the cloud infrastructure. In PhenoMeNal, more than 250 modules were implemented and incorporated into Galaxy. Galaxy also supports more powerful features like programmatic access through a REST API and helper libraries to access the running instance of Galaxy [66].

Jupyter, which started its history as the “IPython notebook”, is the most popular among tools commonly referred to as “executable notebooks” or “computational notebooks” [67]. Jupyter lets users combine executable code with results from code executions such as text, tables and figures. Usually Jupyter notebooks are enriched with extended information that explain what the code does. As a result, they are often used for training material and for tutorials. Computational notebooks can also to some extent be used as a way to document code executions and to make executions more reproducible [68].

Luigi is a Python programming library that was originally developed by the company Spotify. It manages pipelines of computations primarily on Big Data systems such as Hadoop and Apache Spark but also supports local execution [67,68]. Luigi is a very flexible library that facilitates building complex pipelines of batch jobs handling dependency resolution, workflow management and visualization. Similarly, Pachyderm allows to process distributed data and to keep track of the data from every stage of the analysis pipeline [20]. With Pachyderm it is possible to track the provenance of results and to accurately reproduce scientific workflows. Luigi and Pachyderm are well suited for complex scientific tasks and easy to use from the python-environment in Jupyter notebooks without additional integration tooling needed.

In PhenoMeNal, we have extended Galaxy, Jupyter, Luigi and Pachyderm in such a way that they can be orchestrated throughout the cloud infrastructure together with the data analysis tools themselves [69]. Six important metabolomics workflows have been fully integrated into PhenoMeNal (Table 2) and more (mzQuality, NMR-BATMAN) are available for testing (Fig. 6) [212].

### Reproducibility

Three strategies are implemented to ensure technical reproducibility using a continuous integration software development framework [25].

- Infrastructure testing: Procedures were implemented to ensure that each individual component (e.g. the deployment process of software containers, resource management, APIs / ABIs) within the infrastructure is interacting correctly with the other components.
- Container testing: Verification that tools, which are packaged into software containers, build and run correctly in the infrastructure. Dependencies within one container and across several interdependent containers are tested.
- Data testing: The output of tools, which process demonstration data, is checked against a data set that is known to contain the expected result. This is being done for both individual tools and for several tools running in a workflow using the workflow testing tool for Galaxy called wft4galaxy [75].

### Standardization

PhenoMeNal has implemented several dedicated Galaxy modules that directly retrieve and store ISA-Tab data set descriptors from and to MetaboLights, and can convert between other formats. Native Galaxy composite data types to support ISA-Tab and ISA-JSON have also been integrated, building upon the ISA API [22,27]. The ISA data type allows for the upload of an ISA-Tab archive (a zip file containing the ISA set of files and raw data when available), which then is displayed to the users as a single Galaxy history data set. The integrated Galaxy modules include a MetaboLights downloader and uploader (for ingestion and submission), modules to explore study metadata through queries on study factors, ISA-Tab “slicing” where queries are used to select subsets of data files of interest, as well as format conversion (export to ISA-JSON and W4M) and study metadata validation (Table 1).

The ISAcreate module enables the creation of ISA-compliant archives for deposition to repositories such as MetaboLights. The tool presents users with a graphical user interface (GUI) in which to specify study design information such as a treatment plan, sampling and assay plans, as well as QA/QC plans, critical for quality control. During the specification of these plans, the GUI enables semantic markup through the selection of terms chosen from multiple community-based, open ontologies for describing the different components, namely: UBERON ontology for anatomical parts, OBI for experimental protocols [76], MSIO for metabolomics-specific terms and quality control terminology developed by the PhenoMenal project [77,78], DUO for consent and data use terms [79] thereby addressing essential ethical requirements, and STATO for statistical terms (http://www.stato-ontology.org). Based on the combination of the treatment, sampling and assay, and QA/QC plans, the ISA API calculates the experimental graph relationships between subjects, samples, and data files, prospectively. The resulting output is made available as an ISA-Tab history item in Galaxy.

PhenoMeNal also advanced the specification of the nmrML standard data format [75] and contributed a dedicated composite data type for nmrML to Galaxy. nmrML is used extensively throughout the NMR 1D workflow and conversion from raw format into nmrML is supported via dedicated Galaxy modules. Throughout the entire analysis pipeline, modules of computational workflows were designed to accept standard formats such as mzML, XML or CSV whenever possible. Furthermore, the e-infrastructure has been designed in such a way that standardized APIs/ABIs are being used for the programmatic interfaces as well as for deploying services. Modern and standardized programming, scripting and meta languages were selected such as Go, HCL, Python, Shell, XML and YAML that are widely used in cloud computing.

### Reusability

In an ongoing effort, PhenoMeNal is actively advancing the FAIR criteria for good data management and stewardship [80] to be applied not only to data, but also to software tools and computational workflows (Table 3).

**Table 3:**
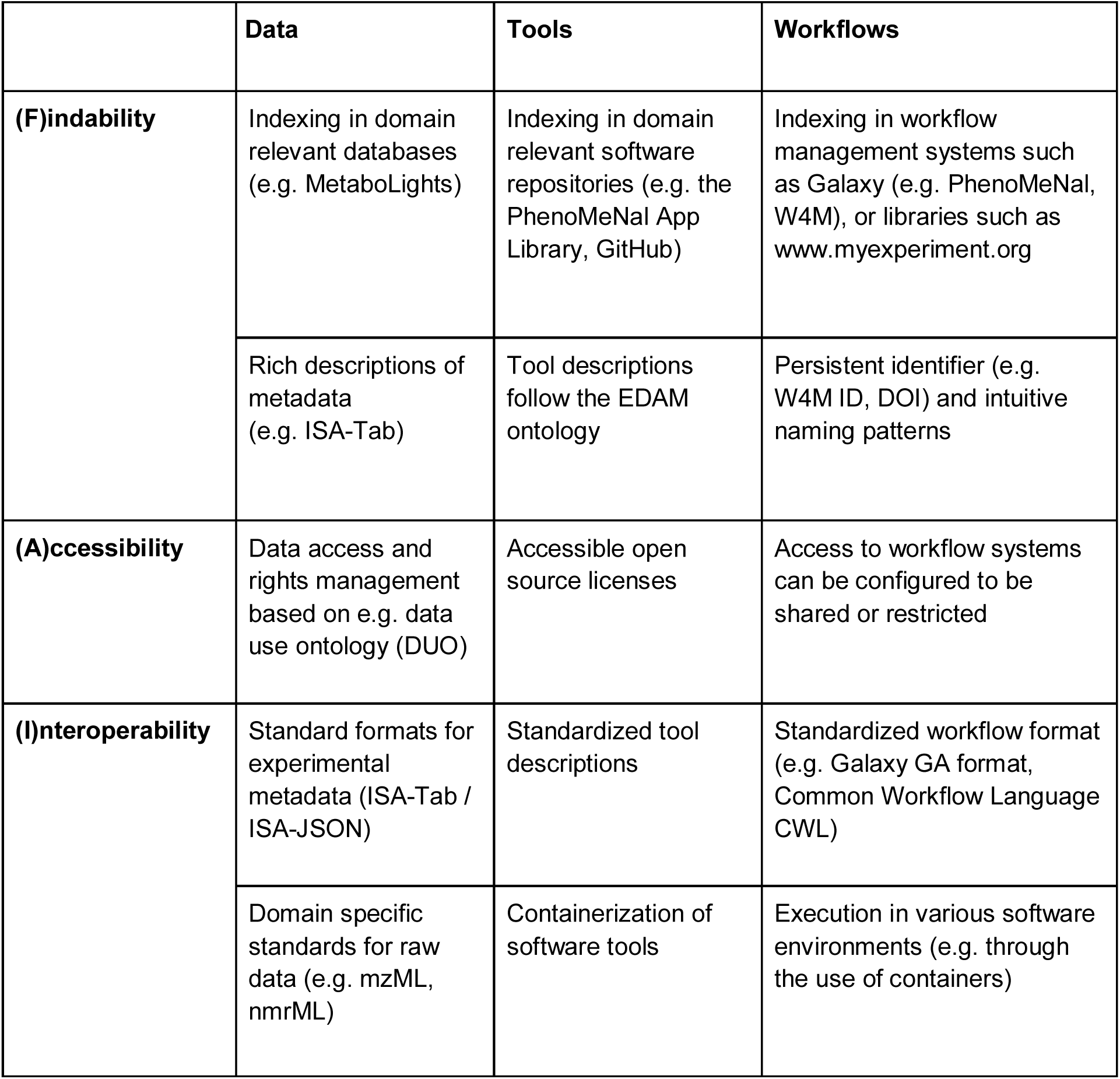
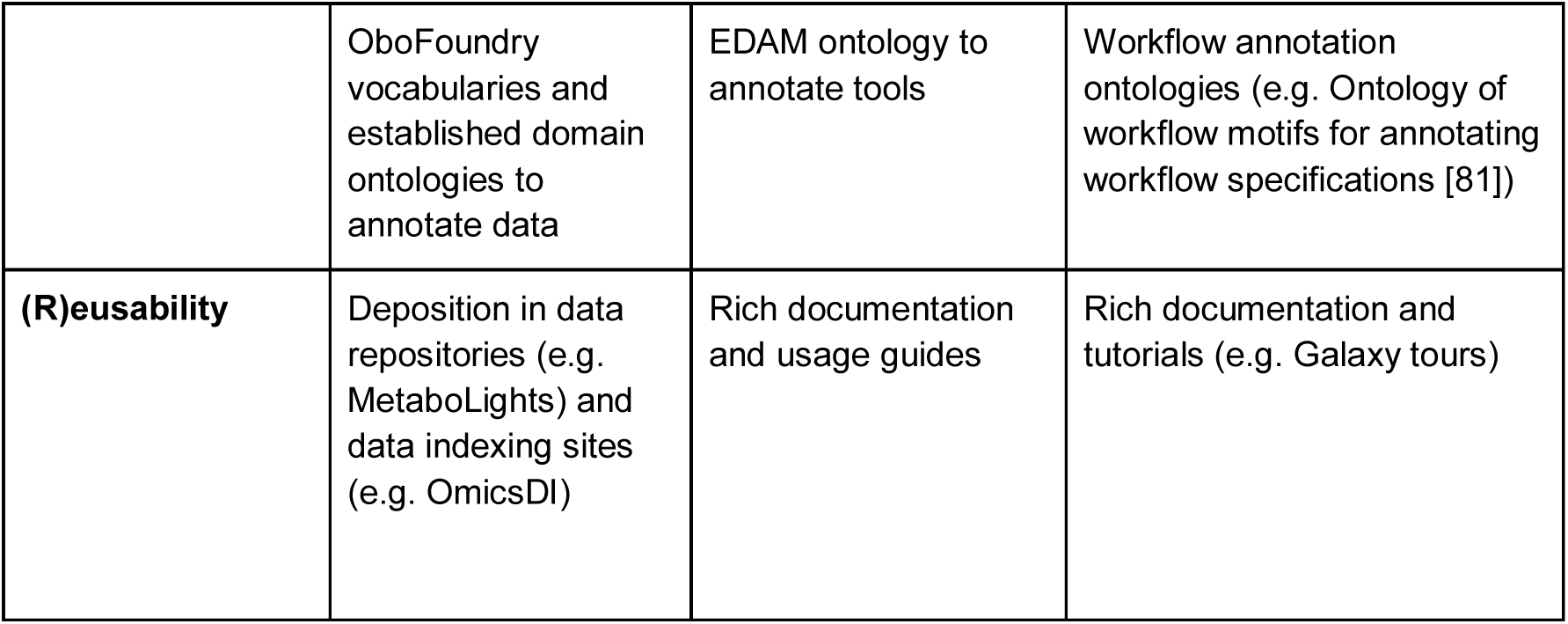
Overview of the most important FAIR criteria and implementations suggested for PhenoMeNal data, tools and workflows.

## Privacy

PhenoMeNal supports fully anonymized data, which cannot be traced back to individuals in any way [29]. This is done by irreversibly removing metadata associated with each individual and any identifier data may be encoded or hashed. It should not be possible, in a reasonable way, to map back to patient identifiers from the data. In this case, the data is generally free from constraints associated with individual patient consent. However, care must be taken when combining with data from other sources not to unexpectedly allow identification of individuals (such as those with very rare diseases or inborn errors of metabolism). Except for such extreme cases, conventional metabolomics data is considered to be non-identifiable. The primary means of achieving this is to require users to only upload fully anonymized data if processing is in a public cloud.

PhenoMeNal follows the guidelines of the European Union and treats pseudonymized data as identifiable. Pseudonymized data are anonymous to the investigator, but allow trusted third parties to link them back to identifiable individuals through mapping such as hash or code [28]. In these cases, e.g. in a hospital environment, users must deploy PhenoMeNal within a private cloud or cluster behind their institution’s firewall. This is consistent with “bringing the compute to the data”, avoiding the need to distribute potentially very large and sensitive data sets across networks of limited bandwidth and unknown security. It is important that ELSI considerations are at the heart of the system, allowing scientists to maintain public confidence that their data are being treated according to their wishes.

PhenoMeNal is fully implementing ELSI and GDPR and has implemented both ethical and technical frameworks to regulate and secure the use of private or sensitive data [28,29]. Thus, patients must have given individual consent to the use of their samples for defined metabolomics research purposes. This information in the European Union is protected under the GDPR^1^. Moreover, in PhenoMeNal, metabolite profile data from animal studies are also treated according to ELSI considerations. In the United Kingdom and Germany, stringent regulations on the use of animals are in place. These legislation rules exceed the requirements of the EU Directive for the protection of animals used for scientific purposes. PhenoMeNal further supports the use of combining data and metadata within an ELSI compliant framework [29] and follows an example of an ELSI compliant architecture, the European Genome Phenome Archive (EGA) [82]. PhenoMeNal follows the ELIXIR policy on privacy and has designed a technical secure environment to process data [26].

### Security

Open source tools are used throughout the entire e-infrastructure and this promotes community efforts to discover and resolve bugs and security issues. The container build process is steered by the key service Jenkins, which continuously builds the containers and generates reports [25]. On success, authentication container images are pushed to the PhenoMeNal container registry which is publicly available but read only. Cloud provider credentials are not stored in the cloud, but only on the deployer host. The Kubernetes cluster running the continuous integration service Jenkins and the container registry, as well as the portal, runs on a CoreOS container, which is a self-updatable, cluster-aware system with most portions being read-only. It reboots nodes sequentially to avoid lack of availability.

KubeNow is a key component that initializes the cloud infrastructure and configures access to it via Cloudflare (http://cloudflare.com/), providing dynamic DNS services and encryption for all communication with services inside the CRE. The flexible implementation of PhenoMeNal allows the user to decide to not use CloudFlare in which case encryption is disabled. KubeAdm, which manages the setup of Kubernetes, is not reachable at runtime by default. The only way to access it is by having access to the private key stored on the computer on which it was launched.

PhenoMeNal only allows access to standard ports (ssh, http, https and port 44 for the Galaxy Downloader) and implements a cloud-specific firewall for all supported cloud providers. Microservices are designed to be launched on-demand and terminated after completed analysis. The deployment uses a base image to speed up provisioning. The latest incremental security patches are applied to the image on startup. Images are built on a daily basis and tested for deployment, to avoid security patches from introducing any abnormality in the deployment process. All virtual machines accept only SSH keys, no passwords are allowed. For long-running services (e.g. Galaxy, Jupyter) the startup script checks and rejects weak application passwords on launch.

### User Resources

A great deal of user resources exist for both PhenoMeNal users and developers, in the form of documentation, tutorials and training videos. The PhenoMeNal Wiki (https://github.com/phnmnl/phenomenal-h2020/wiki) contains detailed documentation on all aspects of PhenoMeNal, including general user guides, workflow and tool tutorials, developer documentation and general information on topics such as security and the e-infrastructure landscape. The PhenoMeNal portal (https://portal.phenomenal-h2020.eu/help) contains help pages generated from the Wiki, which are categorized as User Documentation, Developer Documentation and Workflow Tutorials. Interactive Galaxy tours are directly integrated in Galaxy (https://public.phenomenal-h2020.eu/tours). Training videos are available at the project’s YouTube page (https://www.youtube.com/channel/UCXGAvsVNQk-aUpckjRC8Ang).

## Author contributions

KP and JB contributed equally to the writing of the draft of the manuscript. CS conceived, designed and coordinated the project. PM was the technical lead of the project. The consortium members JB, MCap, MCas, PdA, TMDE, RG, AG-B, UG, KH, SH, DJo, FJ, KK, NK, PEK, AL, MC, PM, SN, COD, KP, LP, MEP, MACR, PR-S, PR-M, AR, RR, CR, MvR, NS, RMS, S-AS, DS, OS, VS, EAT, MT, TH, MvV, MRV, RJMW, GZ, CS contributed to the overall project, including tool development. SB, CF, EH, SH, MI, DJa, BK, IK, KK, PEK, SL, CL, JAN, JTMP, AP, LP, RR were additionally involved with external tool development for the project. All authors contributed to, read and approved the final manuscript.

## Availability of supporting source code and requirements

Project name: PhenoMeNal

Project home page: http://phenomenal-h2020.eu

Operating system(s): Platform independent

Programming language: Go, HCL, Java, JavaScript, Python, R, Shell, XML, YAML

Other requirements: Linux, Docker, Kubernetes, Terraform, Ansible, Helm

License: MIT license for all code written by the PhenoMeNal project. Individual, Open Source Foundation approved licenses for all containerized tools.

## Supporting data

MTBLS1 (NMR1D), MTBLS404 (Uni- and multivariate statistics), MTBLS412 (Fluxomics), MTBLS520 (Eco-Metabolomics), MTBLS558 (MetFrag), available at https://www.ebi.ac.uk/metabolights.

## Declarations

Human-derived samples in the data sets MTBLS404 and MTBLS412 were processed according to ELSI guidelines.

## Author contact addresses

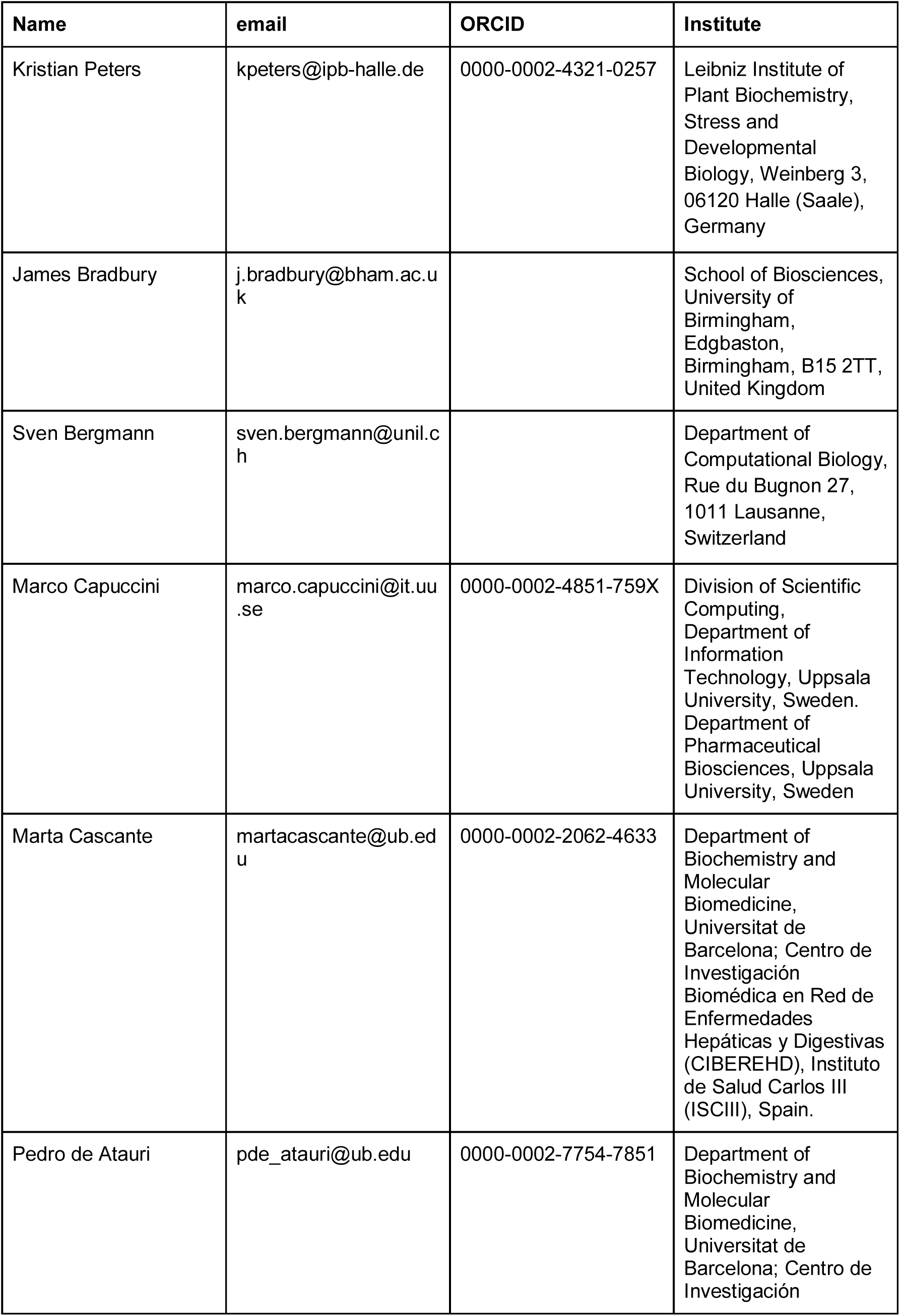
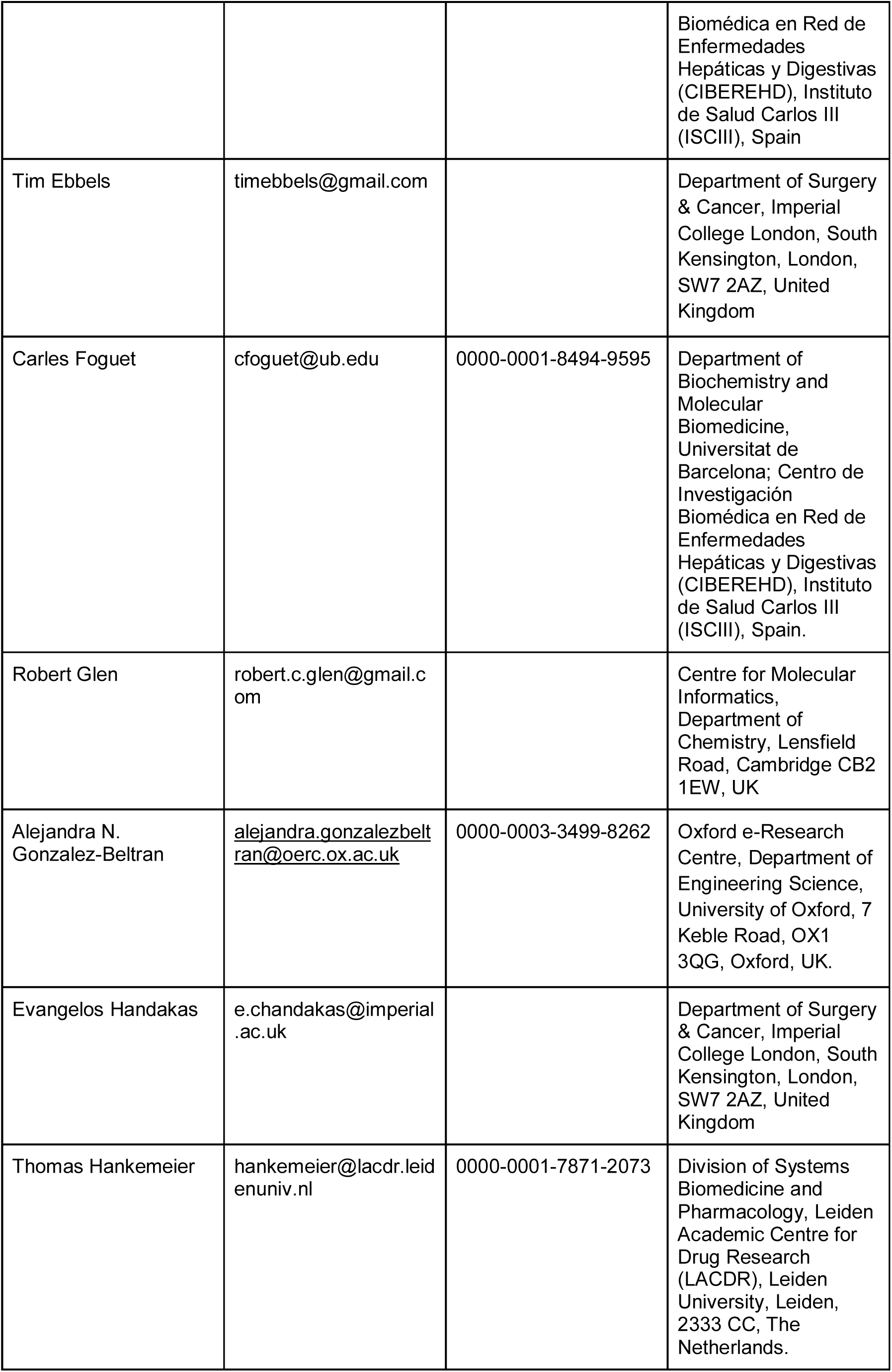
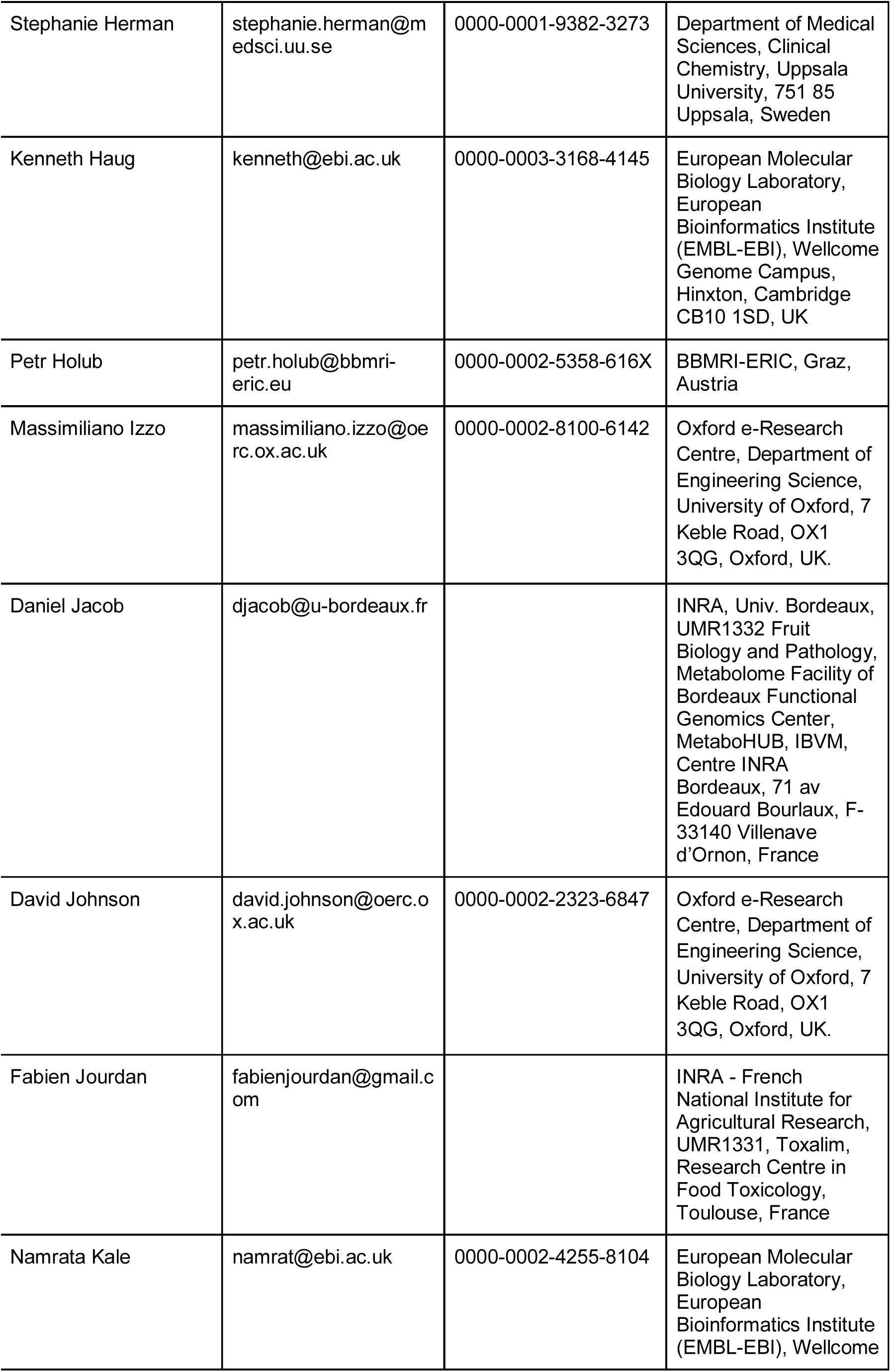
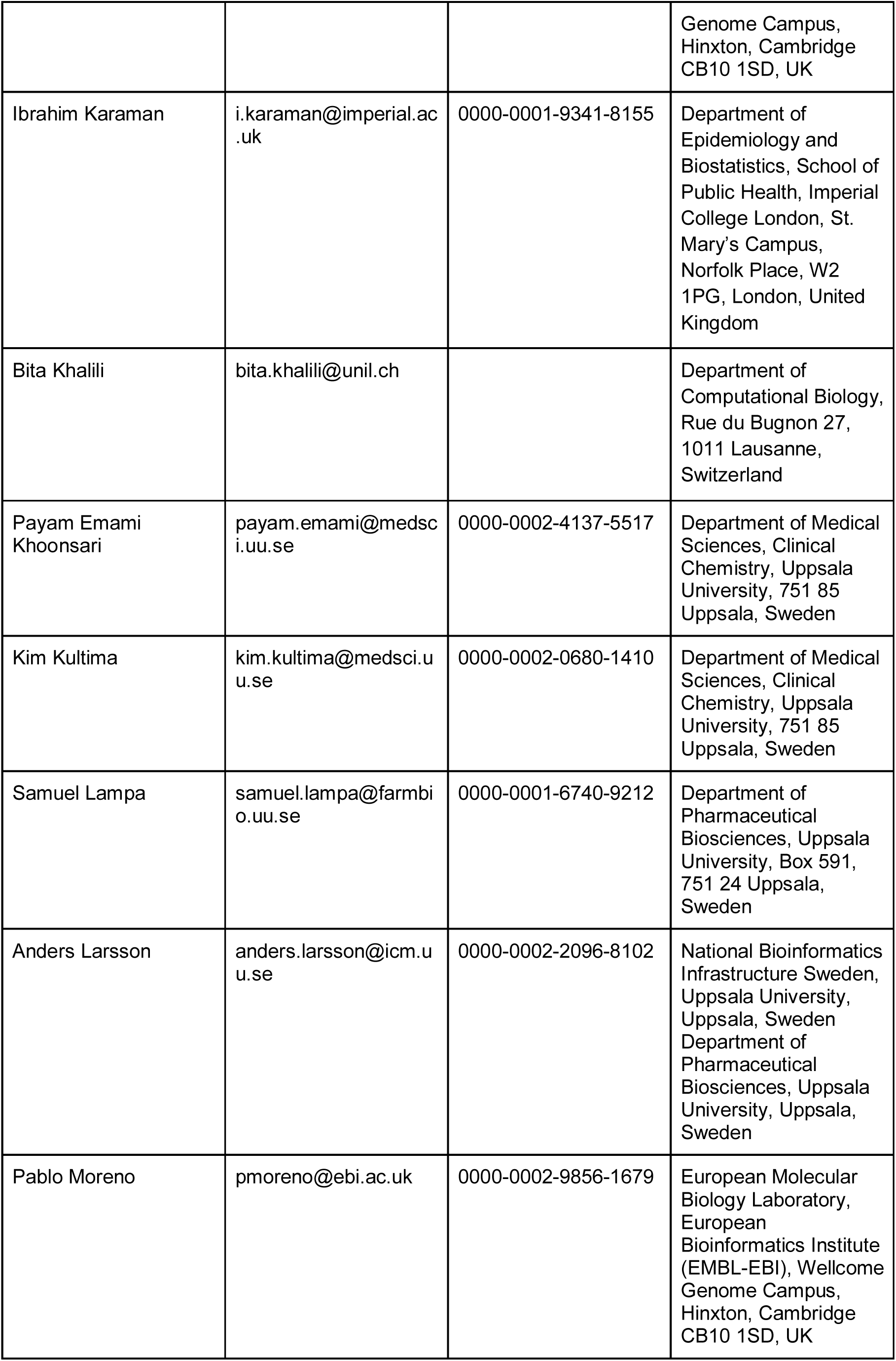
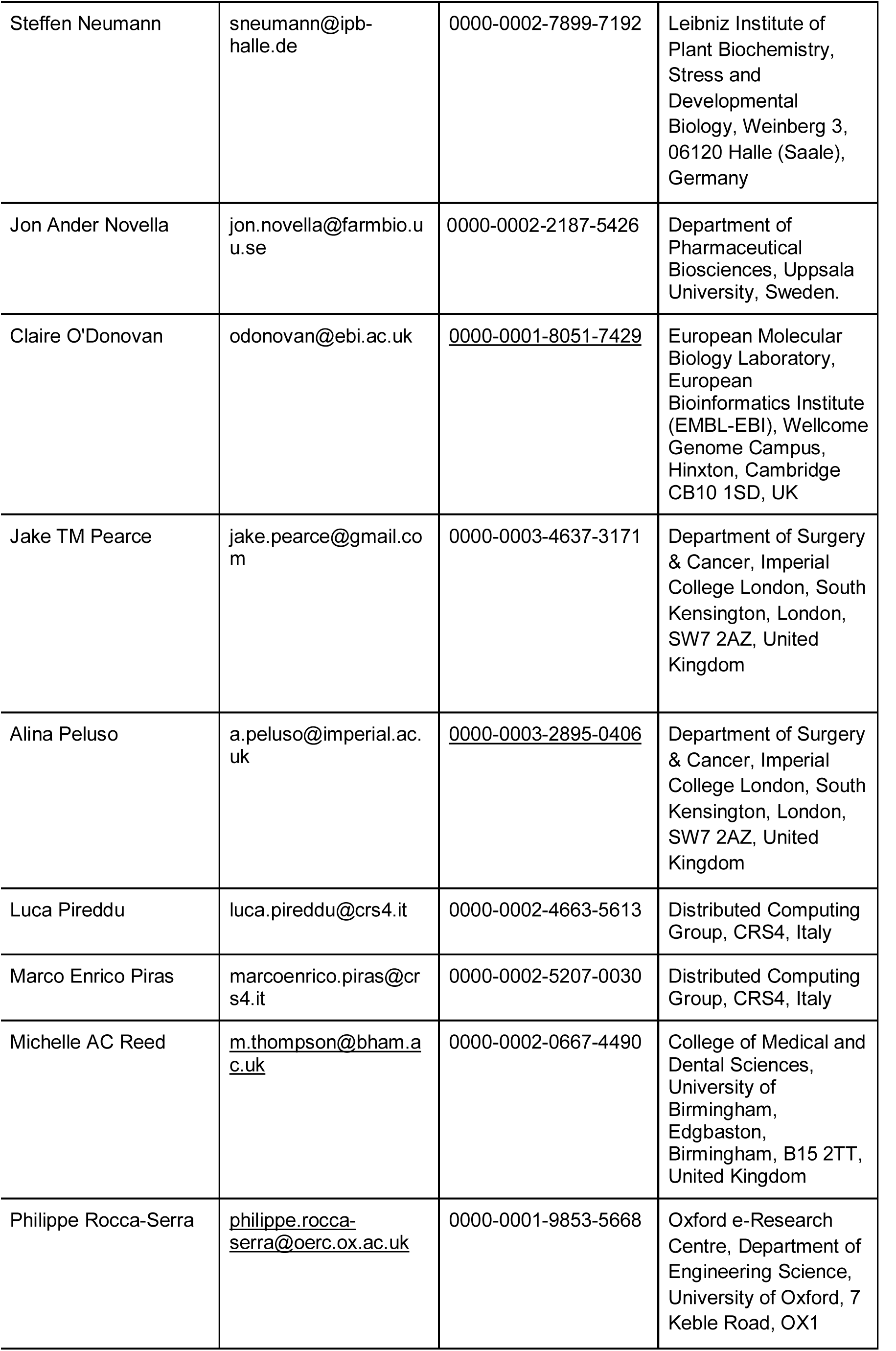
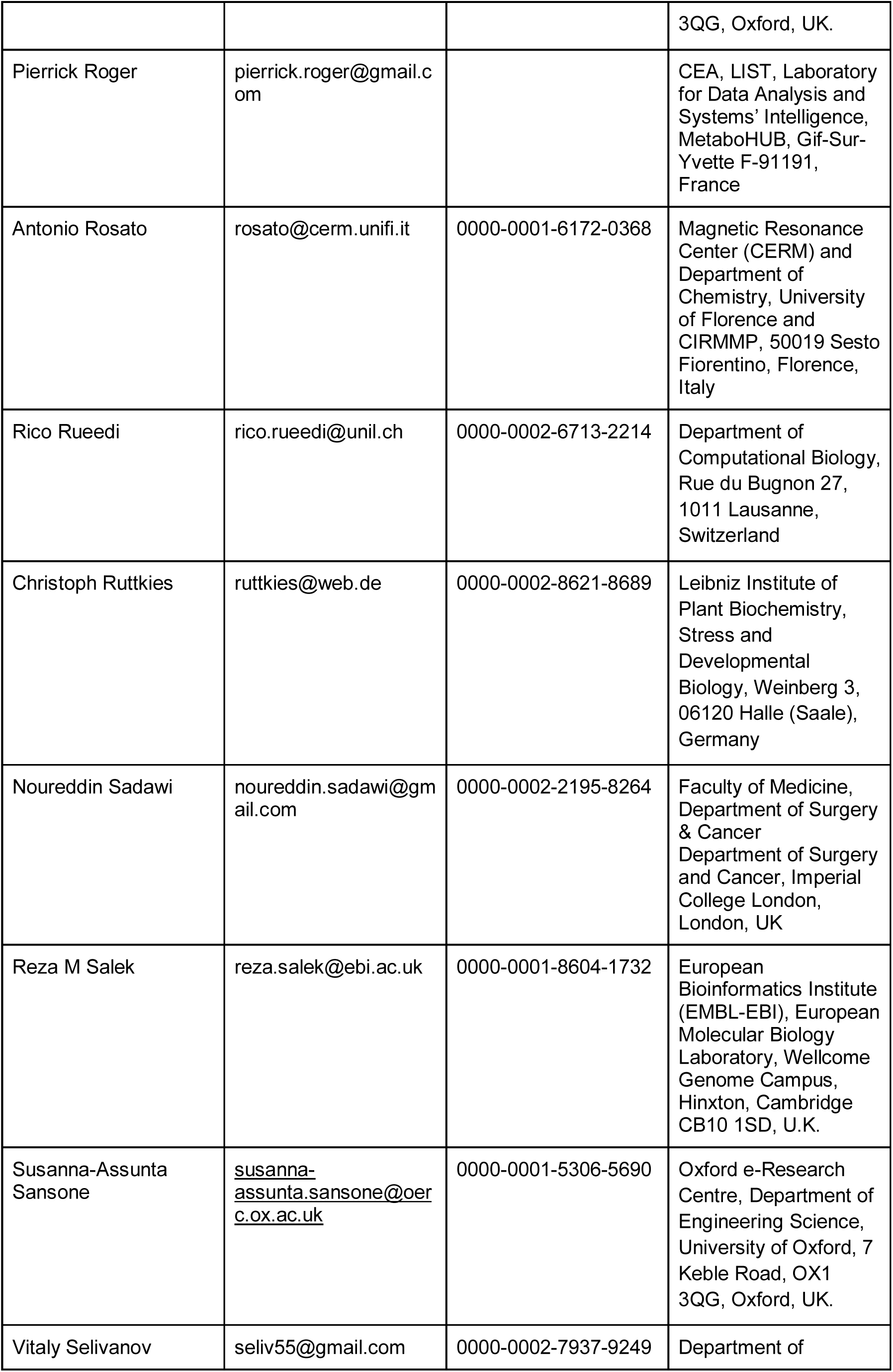
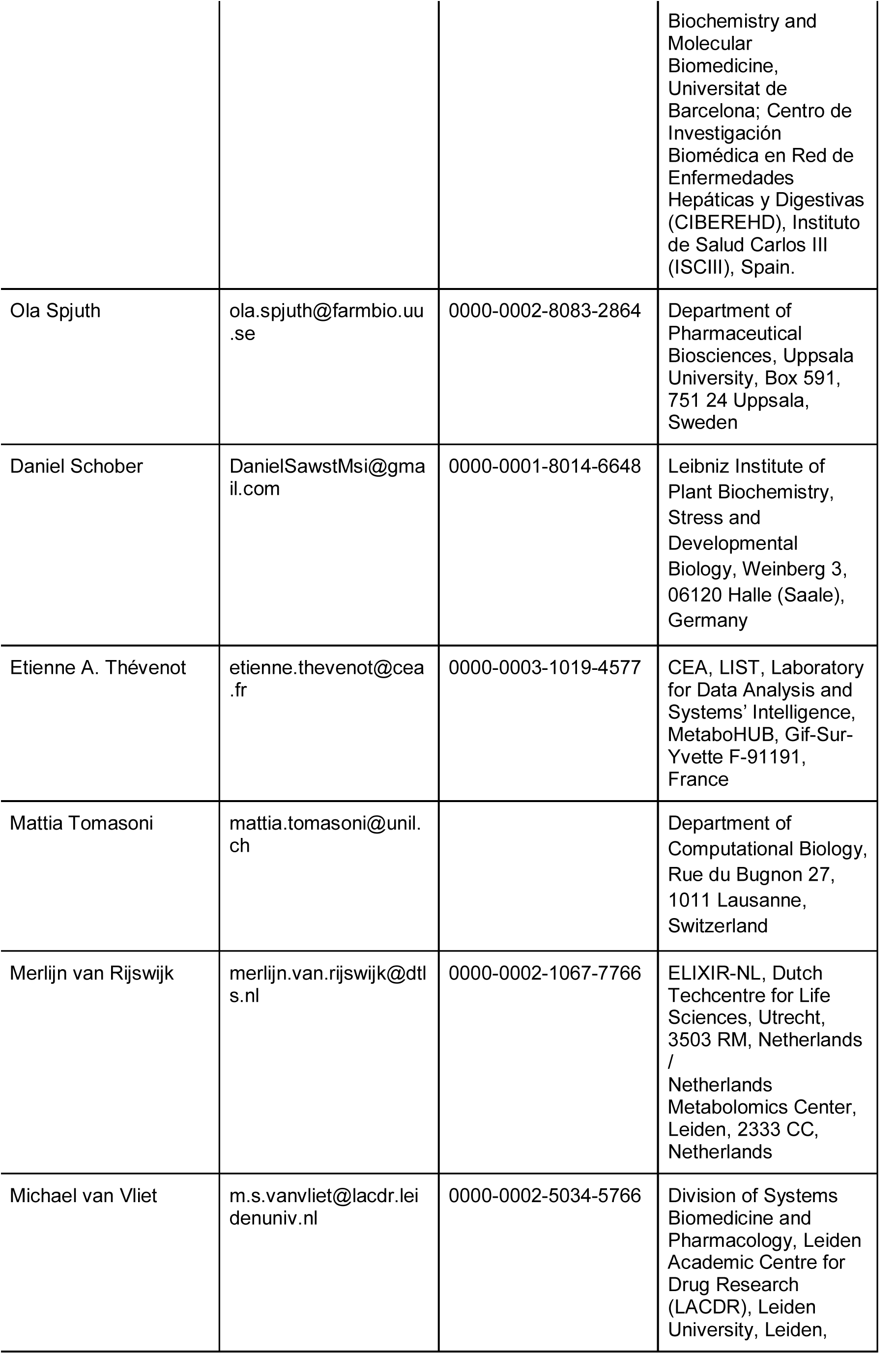
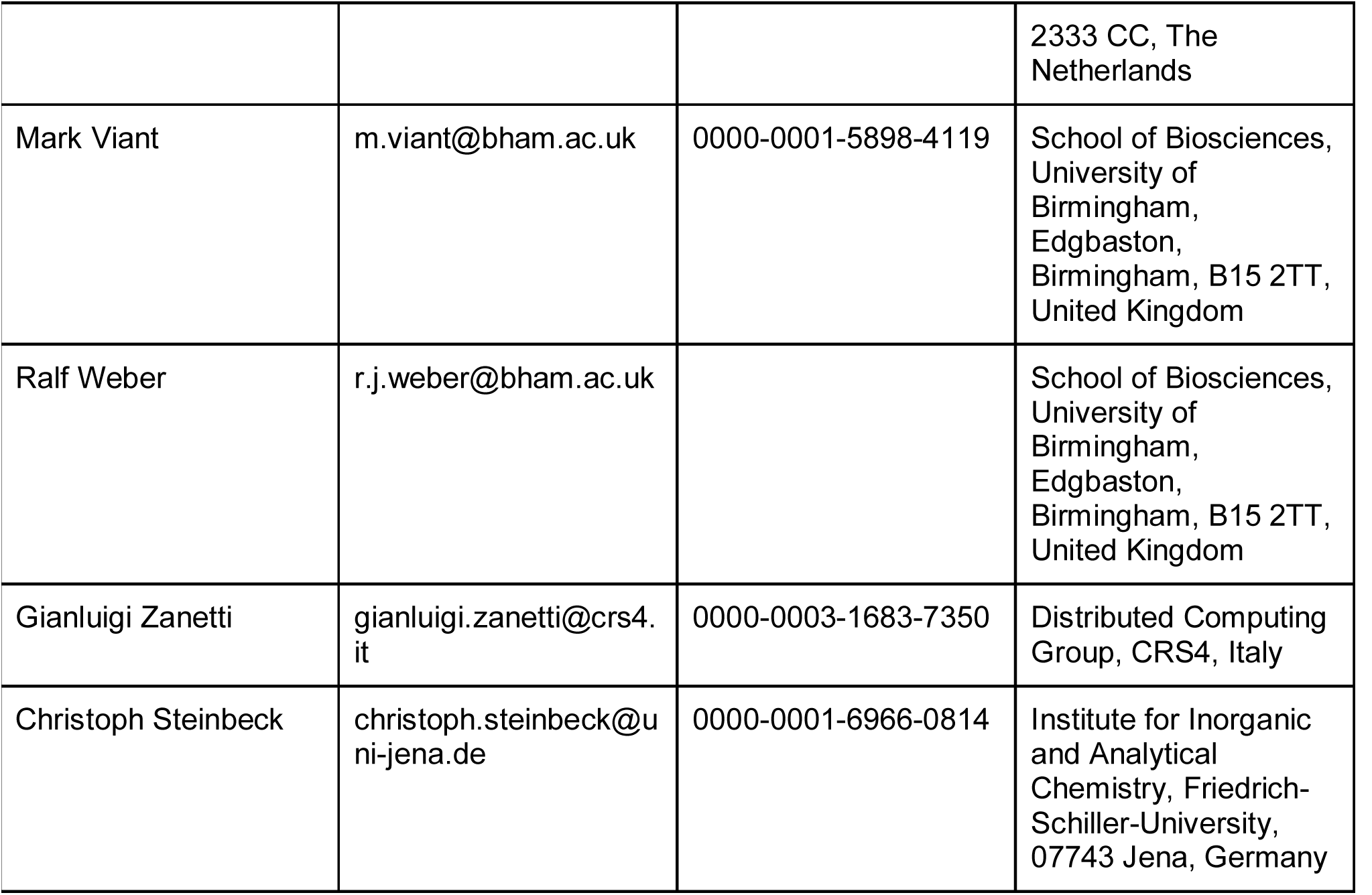

## Footnotes

1 Regulation (EU), 2016/679 of the European Parliament and of the Council of 27 April 2016 on the protection of natural persons with regard to the processing of personal data and on the free movement of such data, and repealing Directive 95/46/EC, 2016, General Data Protection Regulation

